# The anti-HIV Drug Nelfinavir Mesylate (Viracept) is a Potent Inhibitor of Cell Fusion Caused by the SARS-CoV-2 Spike (S) Glycoprotein Warranting further Evaluation as an Antiviral against COVID-19 infections

**DOI:** 10.1101/2020.04.24.060376

**Authors:** Farhana Musarrat, Vladimir Chouljenko, Rafiq Nabi, Achyut Dahal, Seetharama D. Jois, Konstantin G. Kousoulas

## Abstract

Coronaviruses belong to a group of enveloped, positive-single stranded RNA viruses that are known to cause severe respiratory distress in animals and humans. The current SARS coronavirus-2 (SARS CoV-2) pandemic has caused more than 2,000,000 infections globally and nearly 200,000 deaths. Coronaviruses enter susceptible cells via fusion of the viral envelope with the plasma membrane and/or via fusion of the viral envelope with endosomal membranes after endocytosis of the virus into endosomes. Previous results with SARS and MERS CoV have shown that the Spike (S) glycoprotein is a major determinant of virus infectivity and immunogenicity. Herein, we show that expression of SARS CoV-2 S (S-n) glycoprotein after transient transfection of African green monkey kidney (Vero) cells caused extensive cell fusion in comparison to limited cell fusion caused by the SARS S (S-o) glycoprotein. S-n expression was detected intracellularly and on transfected Vero cell surfaces and caused the formation of very large multinucleated cells (syncytia) by 48 hours post transfection. These results are in agreement with published pathology observations of extensive syncytial formation in lung tissues of COVID-19 patients. This differential S-n versus S-o-mediated cell fusion suggests that SARS-CoV-2 is able to spread from cell-to-cell much more efficiently than SARS effectively avoiding extracellular spaces and neutralizing antibodies. A systematic screening of several drugs for ability to inhibit S-n and S-o cell fusion revealed that the FDA approved HIV-protease inhibitor, nelfinavir mesylate (Viracept) drastically inhibited S-n and S-o-mediated cell fusion in a dose-dependent manner. Complete inhibition of cell fusion was observed at a 10 micromolar concentration. Computational modeling and *in silico* docking experiments suggested the possibility that nelfinavir may bind inside the S trimer structure, proximal to the S2 amino terminus directly inhibiting S-n and S-o-mediated membrane fusion. Also, it is possible that nelfinavir mesylate acts on cellular processes to inhibit S proteolytic processing. These results warrant further investigations of the potential of nelfinavir mesylate as an antiviral drug, especially at early times after SARS-CoV-2 symptoms appear.

## Introduction

The severe acute respiratory syndrome coronavirus (SARS-CoV-2) is currently associated with a global pandemic causing coronavirus disease, first noted in December of 2019 in the Wuhan province of China. The resultant disease is termed COVID-19 (coronavirus disease-2019) and is characterized by acute respiratory disease and pneumonia. COVID-19 has infected nearly 2 million people and caused nearly 200,000 deaths worldwide with a predilection of older people and/or people having other health issues including, hypertension, diabetes, obesity and other comorbidities (1–3). SARS CoV-2 is the third human coronaviruses that appeared for the first time in the 21st century. One of the other two coronaviruses are SARS, which appeared in November of 2002 in China and caused 8,098 infections world-wide and 774 deaths. SARS was effectively contained because apparently the virus, although causing high degree of mortality in infected individuals, was not effectively transmitted from one person-to-the other (4, 5). The second human coronavirus, Middle East Respiratory Syndrome Coronavirus (MERS-CoV) appeared in 2013 and caused a limited epidemic of few thousand people, but high death rates of approximately 36% predominantly in the Middle East (Saudi Arabia). The primary source of infection was found to be dromedary camels, although the virus was transmitted from person to person in close proximity in hospital settings (6–9).

All coronaviruses specify a Spike (S) glycoprotein, which is embedded in viral envelopes in trimeric forms giving them their characteristic corona structures. The S glycoprotein is a major antigen responsible for both receptor binding and membrane fusion properties (9). ACE-2 has been identified as the cell receptor for SARS (10), and also SARS CoV-2, while other unknown human receptors may be responsible for its wider infectious spread than SARS. Spike is cleaved into two major components S1 and S2 via cellular proteases. Virus entry into cells is mediated after binding of a receptor-binding domain (RBD) located with the S1 ectodomain. Cleavage of the S glycoprotein to produce S1 and S2 proteins is mediated by cellular proteases at the S1/S2 junction as well as at S2’ site located near the S1/S2 proteolytic cleavage. Fusion of the viral envelope with cellular membranes is mediated by the S2 protein that contains a putative fusion peptide region. The mechanism of membrane pore formation that leads to membrane fusion involves the formation of a six-helix bundle fusion core by two heptad repeats (HR1) and HR2 domains found in each S monomer forming the initial pore that results in membrane fusion (11). The cellular serine protease TMPRSS2 has been implicated in priming S2’ cleavage, as well as ACE2 cleavage both required for initiation of the membrane fusion event (12–14). Also, SARS-S can be cleaved by the cellular protease cathepsin L at the low pH of endosomes, thereby exposing the S2 domain of the spike protein for membrane fusion (15–20). Cell-surface expression of S mediates S-induced cell fusion and the formation of syncytia, which is a phenomenon similar to virus entry requiring the presence of ACE2. Virus-induced cell fusion is a mechanism by which the virus can spread from cell-to-cell by a pH-independent mechanism avoiding the extracellular space and potentially evading neutralizing antibody (21, 22). It has been demonstrated that the RBD domain of SARS CoV-2 S (S-n) has a higher binding affinity for the ACE2 receptor than that of SARS S (S-o), while the S2 proteins of these two viruses are nearly 90% identical (3, 22, 23).

Nelfinavir mesylate was developed as an anti-HIV protease inhibitor (24, 25). In addition to its potent activity against the HIV protease, nelfinavir mesylate was found to produce multiple effects on cellular processes including induction of apoptosis and necrosis as well as induction of cell protective mechanisms, including cell cycle retardation and the unfolded protein response (26–28). These nelfinavir mesylate effects have been exploited for anti-cancer purposes (29–31). Previously, it was shown that nelfinavir mesylate inhibited SARS CoV replication in cell culture (32).

Previously, we investigated the structure and function of the SARS S glycoprotein in transient transfection-membrane fusion assays (33, 34). Based on these initial studies, we undertook screening of a number of compounds that may inhibit S-mediated fusion after transient expression in African green monkey kidney cells (Vero). We report herein that the SARS CoV-2 S (S-n) causes extreme S-mediated membrane fusion in comparison to cell fusion caused by transient expression of S-o. Importantly, we report that nelfinavir mesylate inhibited S-mediated fusion at micromolar ranges. *In silico* docking experiments, revealed the possibility that nelfinavir binds to the S2 amino terminus within the S trimer and thus, may direct inhibit the formation of the heptad-repeat complex that caused S-mediated membrane fusion. Based on these results, further research of nelfinavir’s effect in human COVID-19 patients is warranted.

## Materials and Methods

### Cell line

African green monkey kidney (Vero) cells were maintained in Dulbeco Modified Eagle’s Media (DMEM) with 10% fetal bovine serum (FBS) and 2% primocin (Invitrogen, Inc. CA, USA).

### Reagents and Antibodies

Nelfinavir mesylate and DMSO were bought from Sigma, Inc (MO, USA). The primary antibodies used were as follows: mouse anti-myc antibody (Abcam, MA, USA), mouse anti-FLAG antibody (Abcam, MA, USA). Goat anti-mouse antibody conjugated with HRP (Invitrogen, Inc. CA, USA) was used as a secondary antibody. The Vector Nova Red peroxidase (HRP) substrate kit (vector laboratories, CA, USA) was used for imaging. Goat anti-mouse antibody conjugated with alexa fluorophore 647 and goat anti-rabbit antibody conjugated with alexa fluore 488 (Invitrogen, Inc. CA, USA) were used for immuno-fluorescence (IFA) assay.

### Construction of Recombinant Spike Proteins

The SARS (S-o) and SARS-2 (S-n) Spike expression plasmids used in the present study were constructed in a very similar manner. Both S genes were placed under the control of the human cytomegalovirus immediate early (CMV) promoter and were engineered to contain 3xFLAG and N-myc epitope tags at their amino terminal ends, respectively. These S-o and S-n were cloned in to p3xFLAG-CMV-9 (Sigma) and pCMV3-SP-N-MYC (Sino Biological) parental vector plasmids, respectively. The S1 subunit of the S-n expression construct contained the same amino terminus up to aa 700 (Gly). The N terminal domain of the S2 S-n subunit was engineered to be exactly as in S1 containing the N-myc tag at its amino terminus and encompassing the S2 S-n amino acid sequence aa701-aa1273.

### Transient Transfection Assay

Vero cells were grown on 24 well plates and transiently transfected with either pCMV3-SP-N-MYC (S-n) or p3XFLAG-CMV-S (S-o) using lipofectamine reagent. Approximately, 2microliter of lipofectamine and 0.5 micrograms of plasmid DNA was used for transfection per well of Vero cells. Appropriate controls were also used. Following 48 hours, the plates were examined by phase contrast microscopy for fused cells and images were taken under live conditions, as well as either after formalin or methanol fixation. Cells were stained for FLAG (S-o, anti-FLAG-1:2500) or N-myc (S-n, anti-myc-1:500) with HRP (Vector Nova Red kit) for phase contrast microscopy. Similarly, cells were stained for fluorescent microscopy using anti-mouse antibody conjugated with Alexa fluorophore 647 and anti-rabbit antibody conjugated with alexa fluorophore 488 (1:1000).

### Inhibition S-mediated Cell Fusion by Nelfinavir

Nelfinavir mesylate was dissolved in DMSO at 10mM concentration (stock) and a series of dilutions was made in serum free DMEM. Following transfection, 500 microliter of nelfinavir mesylate solution was added to each well. Vero cells transfected with either S-o or S-n and incubated with the drug for 48 hours at 37°C with 5% CO2. The tissue culture plates were observed for fused cells, and then phase contrast and fluorescent images were taken under either formalin or methanol fixed conditions.

### Computational Methods

Docking of the nelfinavir mesylate to the spike protein of SARS CoV-2 was performed using Autodock (35). Crystal structure of nelfinavir was obtained from the complex of HIV-protease nelfinavir crystal structure from the protein data bank (PDB ID: 2Q64) (36). Structure of the prefusion S protein of 2019-nCoV was reported by Wrapp et al., (2020) (PDB ID: 6VSB)(37). The trimer structure of the spike protein was used for docking as protein structure of the spike protein exists under dynamic condition while binding to the receptor and fusion to host cell. Grid for docking was created on the spike protein structure at particular docking site as the center covering a grid box of 126 Å in X, Y, Z direction from the center of the grid. One grid site was created around protease cleavage site S1/S2 and another covering the HR1 region of the protein in the trimer (**Figure S1, in supporting information**). Docking calculations were performed using Lamarckian genetic algorithm with 150 starting conformations and 10 million energy evaluations. Fifty low energy docked structures were used for final analysis. Structures within 2 kcal/mol from the lowest energy docked structures were represented as final possible docked structures using PyMol software (Schrodinger).

Instruments and Software. Olympus IX71 fluorescent microscope was used for live and phase contrast images using Cellsens software. Zeiss Axio Observer Z1 fluorescent microscope was used for fluorescent images using Zen software.

## Results

### SARS CoV-2 Spike (S-n) is drastically more fusogenic than SARS Spike (S-o)

Virus entry is achieved by S-mediated fusion between the viral envelope and either cellular plasma membranes or endosomal membranes. S-mediated cell fusion is caused by cell-surface expression of S and it is thought to be a surrogate model of both virus entry and cell fusion. Previously, we reported a detailed analysis of the functional domains of the SARS Spike (S) glycoprotein that are important for S-mediated membrane fusion and the formation of multi-nucleated cells (syncytia) including delineation of domains important for synthesis, cell-surface expression and endocytosis from cell surfaces (14, 15). To compare the S-o-mediated fusion versus the S-n, both genes were cloned into the transient expression vectors as codon-optimized genes carrying a 3XFLAG or N-myc epitope tags at their amino termini **(Fig. 1: A, B, E, and F).** In addition, S1 and S2 domains of S-n were cloned independently into the transient expression vector pCMV3, encompassing amino acid domains for S1 (aa16-aa700) and S2 (aa701-aa1273). Both S1 and S2 domains were expressed with a myc epitope tag at their amino termini (Fig. 1, C, D). Vero cells were transfected with the S-n or the S-o expressing plasmids and were detected at 48 hours post transfection (hpt) using anti-myc and anti-FLAG antibodies in conjunction with secondary antibody linked to horseradish peroxidase (see Materials and Methods). Vero cells were also transfected with plasmid vehicle controls or mock-transfected. Expression of both S-n and S-o was readily detected by immunohistochemistry, while there was no signal obtained from the Vero mock-transfected and HRP-stained control cell monolayers. Phase contrast microscopy revealed the presence of extensive syncytial formation in S-n, but not S-o transfected cells, while the remaining monolayer did not appear to exhibit any cellular toxicity **(Fig. 2A)**. Further examination of transfected Vero cells by immunofluorescence staining for cellular tubulin (anti-alpha tubulin antibody), nuclei (DAPI) and anti N-myc and anti-FLAG antibodies followed by anti-mouse fluorescent antibody provided additional support that untransfected monolayers appeared normal, while S-n expression produced large syncytia in contrast to much smaller syncytia formed after S-o transient expression **(Fig. 2B).** Expression of either S1, S2 or S1+S2 domains of S-n was readily detected by immunohistochemistry with the anti-N-myc antibody; however, there was no substantial cell fusion observed at 48 hpt (**Fig. 3**), as well as at later times (not shown).

**Figure 1.**
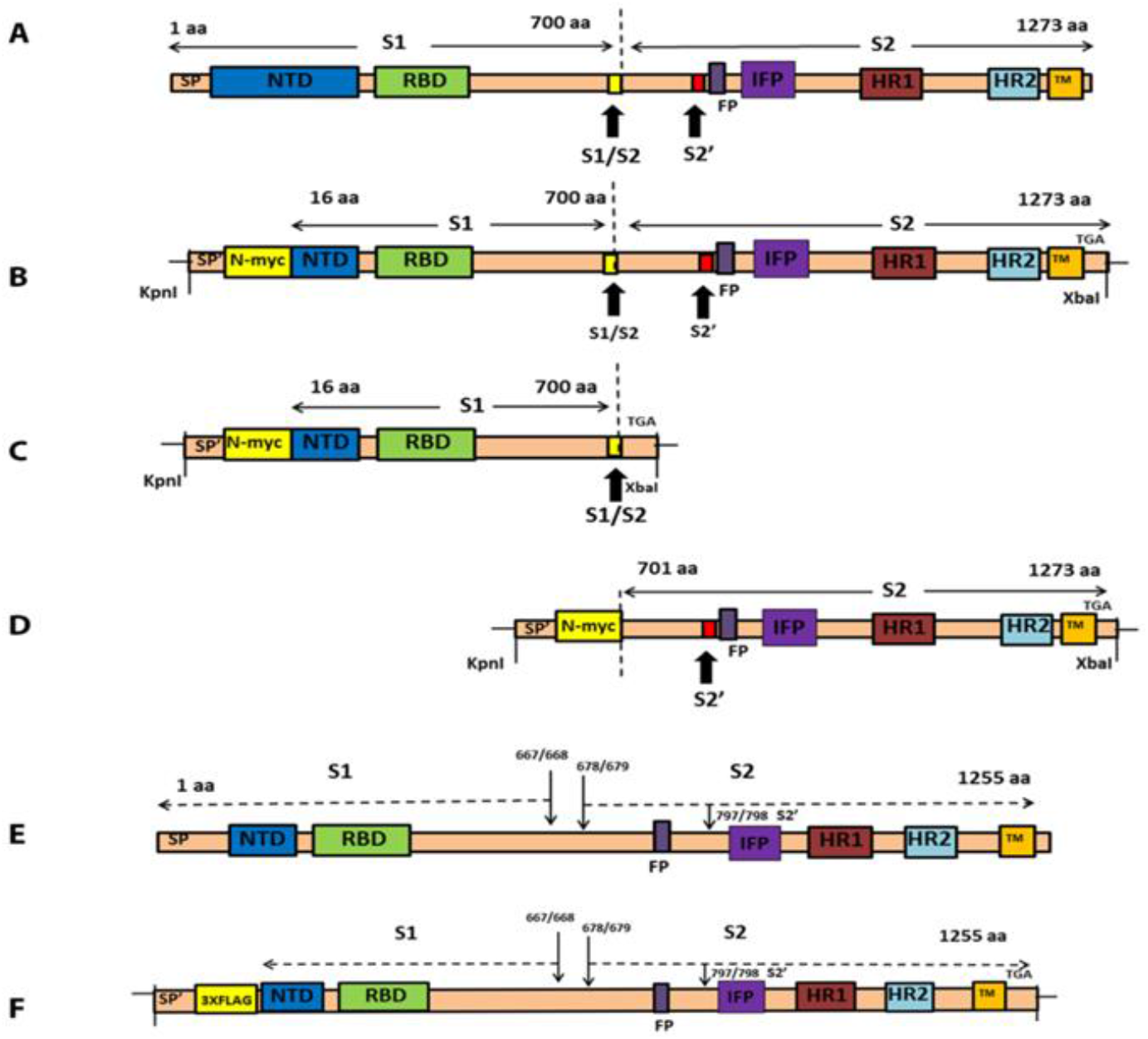
Schematics of spike glycoproteins and recombinant gene constructs. A) Structure of SARS-CoV-2 spike (1273 aa) glycoprotein, showing S1 and S2 domains and the cleavage sites S1/S2 and S2’. B) Structure of pCMV3-SP-N-MYC (S-n). SARS-CoV-2 spike (aa16-aa1273) was cloned into pCMV3-SP-N-MYC at Kpnl and Xbal restriction sites. The N-terminal 15 amino acids were replaced with signal peptide (SP’) and N-myc sequence. C) Structure of pCMV3-S1-N-MYC (S1-n). The S1 domain (aa16 – aa700) was also cloned into pCMV3-SP-N-MYC at Kpnl and Xbal restriction sites. The N-terminal 15 amino acids were replaced with signal peptide (SP’) and N – MYC sequence. D) Structure of pCMV3-S2-N-MYC (S2-n). The S2 domain (aa701– aa1273) was cloned into pCMV3-S2-N-MYC at Kpnl and Xbal restriction sites. The N-terminal contains signal peptide (SP’) and N-myc sequence. E) Structure of SARS spike (aa1255) glycoprotein, showing S1 and S2 domains and the cleavage sites S1/S2 and S2’.F) Structure of p3XFLAG-CMV-S (S-o). SARS spike expression plasmid was constructed as described previously (33, 34), and was similar except 3X FLAG instead of N-myc was used for detection. HR1= heptad repeat 1, HR2= heptad repeat 2, NTD= non translated domain, RBD= receptor binding domain, FP= fusion peptide, SP= SARS signal peptide, SP’= signal peptide

**Figure 2.**
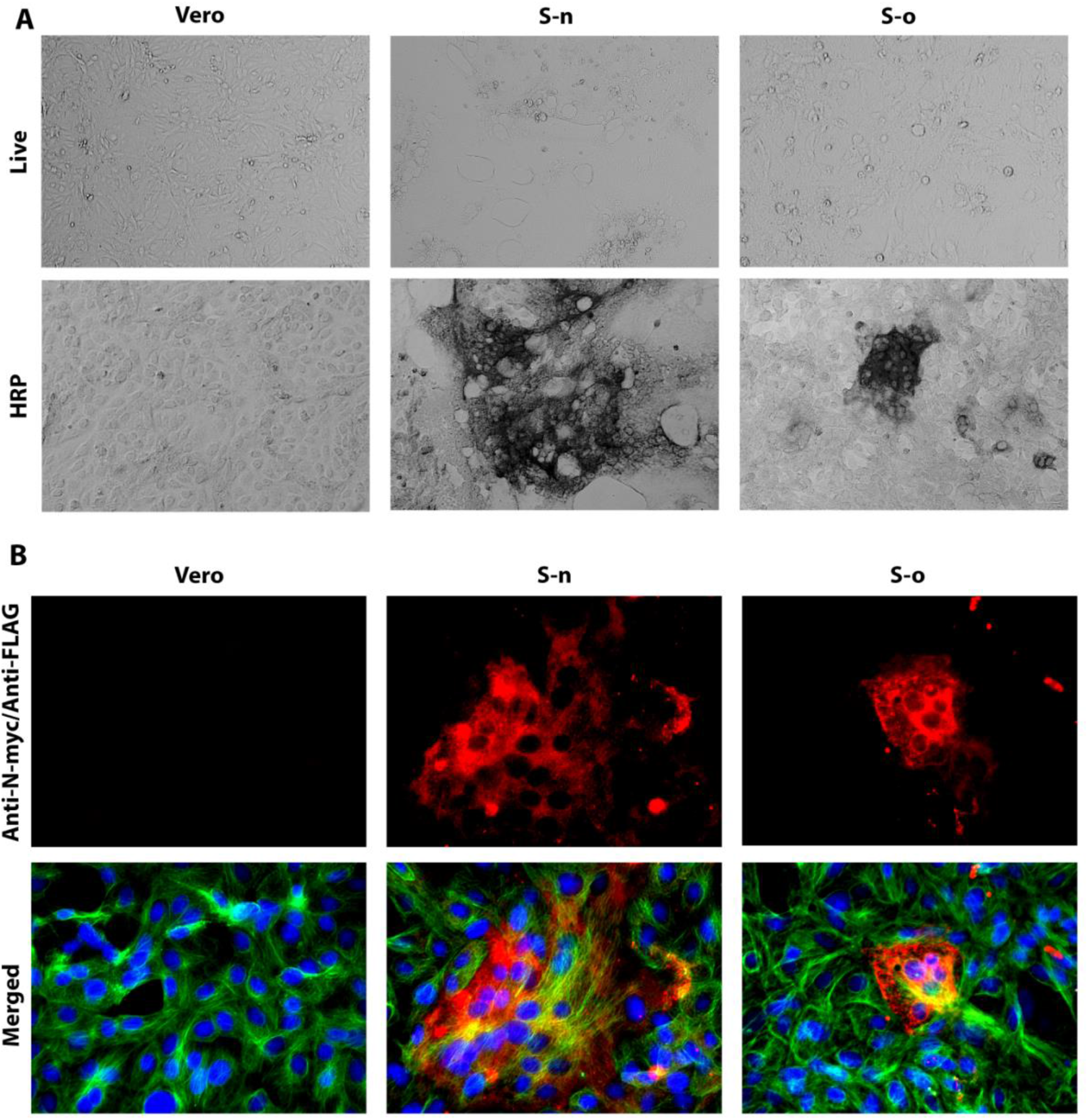
Syncytia formation by S-n and S-o. Vero cells were transfected with plasmid vectors expressing either the S-o or S-n glycoproteins tagged with the 3xFLAG and N-myc epitopes at their amino termini, respectively. S-n and S-o expression was detected with mAbs against the epitope tags at 48 hours post transfection and compared to vehicle containing equivalent amount of lipofectamine. Methanol fixed cells were incubated with mouse anti-N-myc (S-n) (1:500 or 1:50) or mouse anti-FLAG (S-o) (1:2500 or 1:200) antibody and stained with either (A) HRP staining or (B) Alexa fluore 647 conjugated goat anti-mouse secondary antibody (1:1000). Cellular tubulin was stained with rabbit anti-alpha tubulin (abcam; 1:200) and anti-rabbit secondary antibody conjugated with alexa fluore 488. DAPI was used to stain nuclei of cells. Phase contrast images were taken at X10 magnification, whereas the fluorescent images were taken at 40X magnification.

### Nelfinavir significantly inhibits cell-to-cell fusion mediated by S-n and S-o without affecting cell-surface expression

Transiently transfected Vero cells were treated with either DMSO, or a series of dilutions (100uM-0.001 uM) of nelfinavir mesylate. Following 48 hours post transfection, the cells were fixed with methanol and stained for either N-myc (S-n) or FLAG (S-o). Nelfinavir mesylate treatment did not inhibit overall S-n and S-o expression, as evidenced by the efficient expression and detection of both proteins via immunohistochemistry. Both S-n and S-o mediated fusion was significantly inhibited by nelfinavir at a dose dependent manner with complete inhibition observed at the lowest effective concentration of 10 micromolar compared to the untreated control **(Fig. 4: A, B)**. To determine the effect of nelfinavir on the surface expression of spike, we transiently transfected Vero cells with plasmids expressing either the S-o or S-n glycoproteins tagged with the 3xFLAG and N-myc epitopes at their amino termini, respectively and treated these cells with either nelfinavir (10uM) or DMSO for 48 hours at 37°C with 5% CO2. The cells were observed for characteristic syncytia formation, and then fixed with either formalin or methanol to detect surface expression or endogenous expression of the spike glycoprotein following nelfinavir treatment. Although there were significant differences in the number of fused cells (size of syncytia) following drug treatment, no apparent difference was visible in the surface expression of spike compared to total spike expression between Sn and So-transfected cells. These experiments revealed that nelfinavir at concentrations that significantly inhibited cell fusion, did not affect S-n or S-o cell surface expression (**Fig. 5)**.

**Figure 3.**
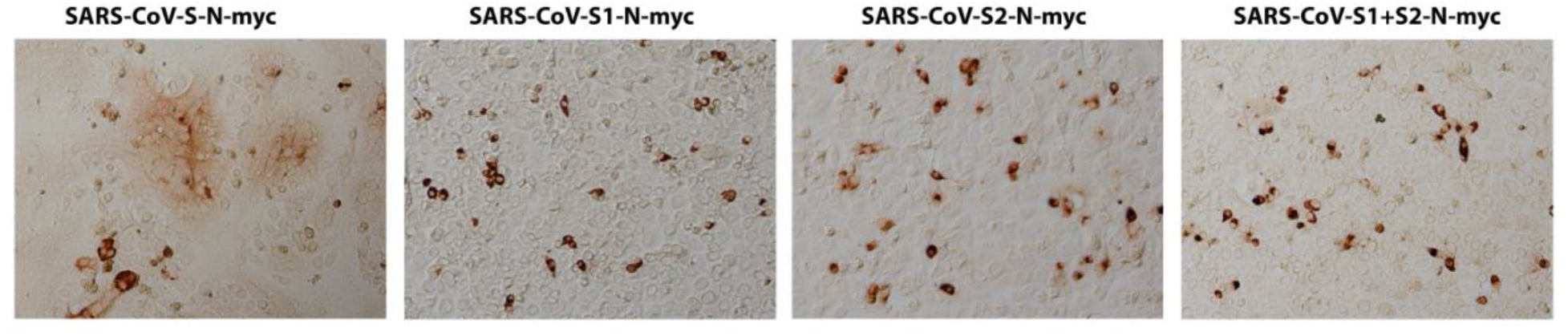
Expression of SARS CoV-2 spike domains. Vero cells were transfected with plasmids expressing either the S1, S2 or S1+S2 domains tagged with the N-myc epitopes at their amino termini. Expression was detected with mAbs against the epitope tags at 48 hours post transfection and compared to vehicle containing equivalent amount of lipofectamine. Methanol fixed cells were incubated with mouse anti-myc antibody and stained with HRP staining followed by goat anti-mouse secondary antibody incubation. Images were taken at 10X magnification.

**Figure 4.**
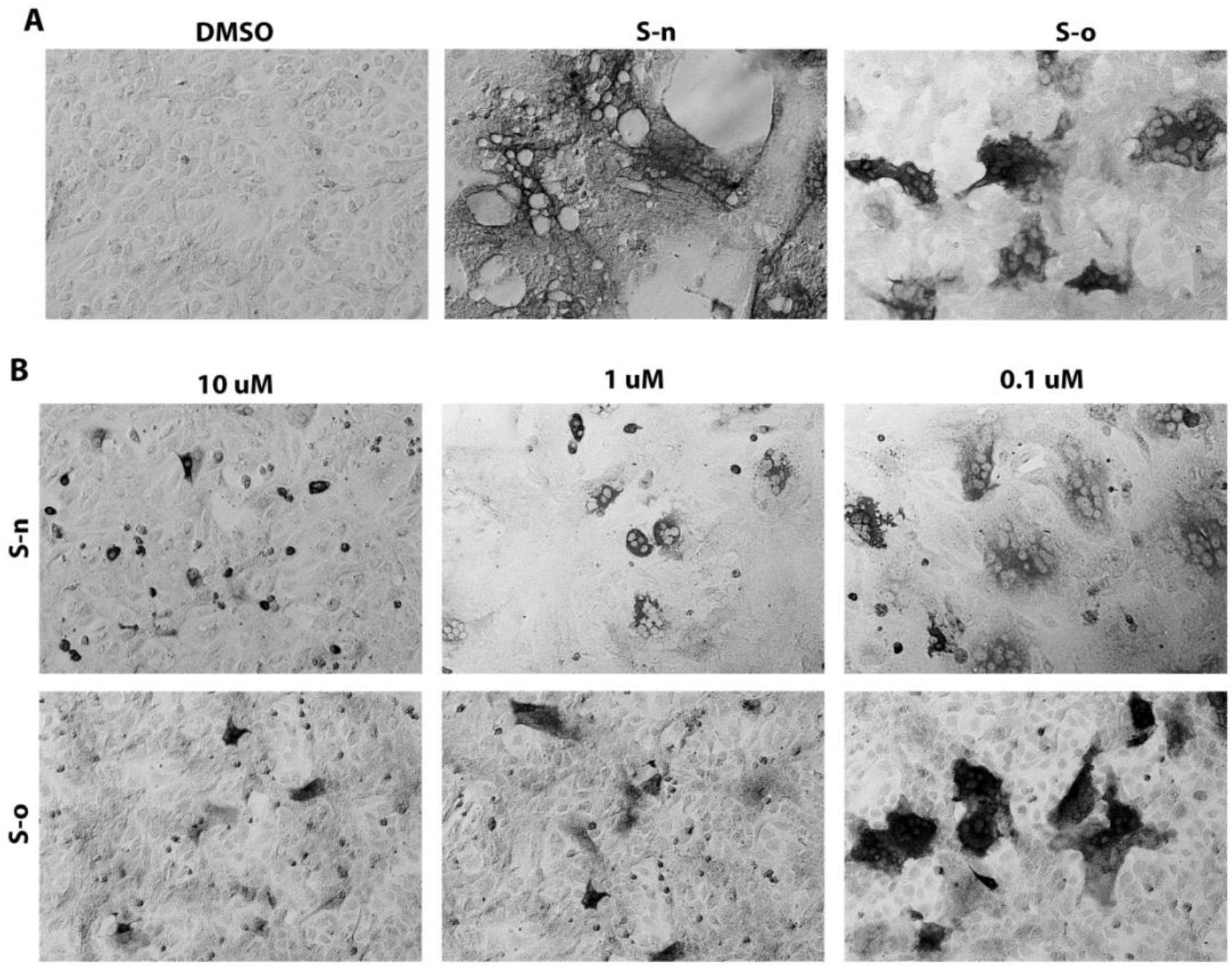
Fusion inhibition by nelfinavir. (A) Vero cells were transfected with plasmids expressing either the S-o or S-n glycoproteins tagged with the 3xFLAG and N-myc epitopes at their amino termini, respectively. S-n and S-o expression was detected with mAbs against the epitope tags at 48 hours post transfection and compared to vehicle containing equivalent amount of DMSO. (B) S-n and S-o glycoproteins were expressed as in A. Nelfinavir was added at the time of transfection at the concentrations indicated. Methanol fixed cells were incubated with mouse anti-N-myc (S-n) or mouse anti-FLAG (S-o) antibody and stained with HRP staining followed by goat anti-mouse secondary antibody incubation. Images were taken at X10 magnification.

**Figure 5.**
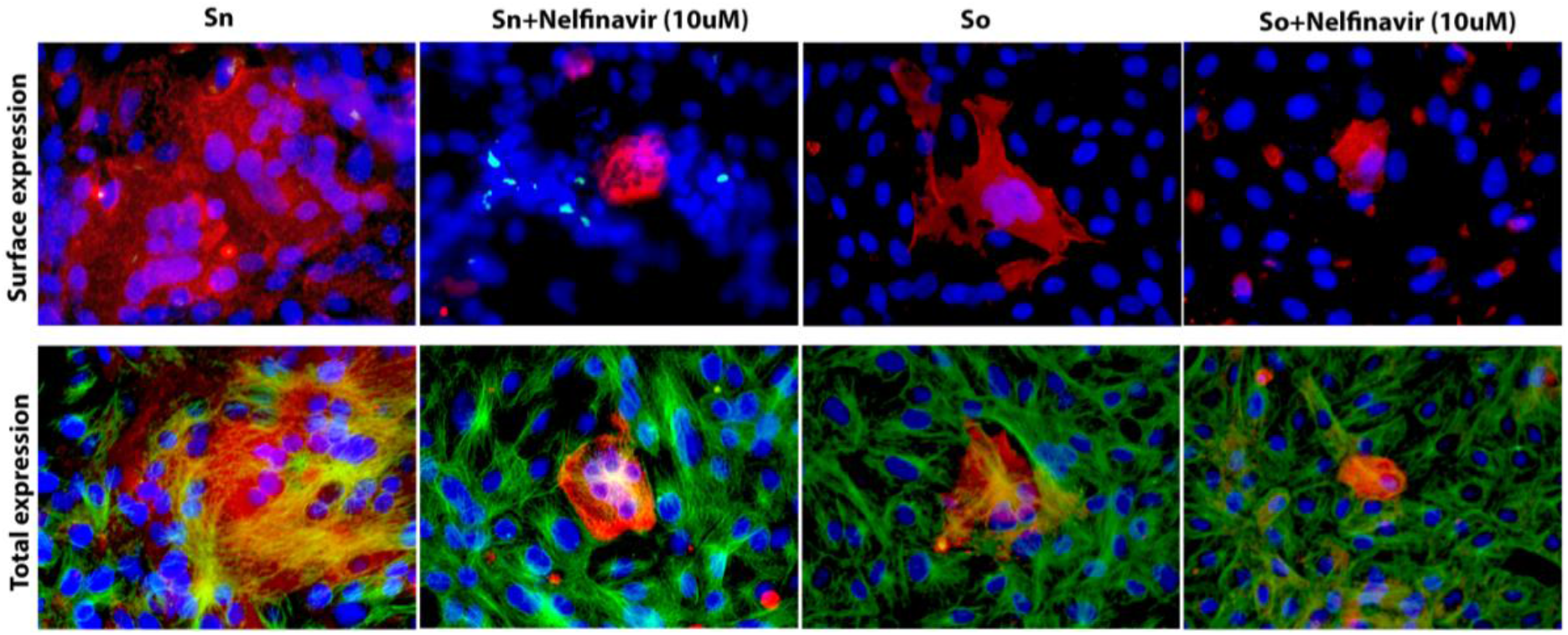
Surface expression of spike glycoproteins. Vero cells were transfected with plasmids expressing either the S-o or S-n glycoproteins tagged with the 3xFLAG and N-myc epitopes at their amino termini, respectively. S-n and S-o expression was detected with mAbs against the epitope tags at 48 hours post transfection and compared to vehicle containing equivalent amount of DMSO. Nelfinavir was added at the time of transfection at the concentrations indicated. Formalin or Methanol fixed cells were incubated with mouse anti-N-myc (S-n) (1:100) or mouse anti-FLAG (S-o) (1:200) antibody and stained with Alexa fluore 647 conjugated goat anti-mouse secondary antibody (1:1000). Cellular tubulin was stained with rabbit anti-alpha tubulin (abcam; 1:200) and anti-rabbit secondary antibody conjugated with alexa fluore 488. DAPI was used to stain nuclei of cells. Fluorescent images were taken at 40X magnification.

### Computation modeling of nelfinavir-S-n potential interactions

*In silico* docking experiments were performed to investigate whether nelfinavir has the potential to directly bind S-n glycoprotein. Two docking grids were created to specifically encompass S1/S2 cleavage site and nearby area and the HR1 region of the trimeric form of S-n (Fig. S1). For the docking grid that was was created to cover the S1/S2 cleavage site, the low energy docked structure of nelfinavir was bound in the pocket between the helices of fusion peptide and HR1 region and lower part of NTD region (Fig. 6A). The docking energy of the nelfinavir bound structure was −9.98 kcal/mol. In the lowest energy docked conformation, the nelfinavir-2019-nCoV complex was stabilized by three hydrogen bonds and hydrophobic interactions. T768 from S protein fusion peptide formed two hydrogen bonds and Q957 of HR1 helix formed one hydrogen bond with nelfinavir. Hydrophobic interaction was dominated by aromatic functional groups of nelfinavir with Tyr313, Leu303 and Q314 side chains alkyl group in the S protein (Fig. 6A). For the docking grid around HR1 helices of the trimer, the lowest energy docked structure was bound near the helices of HR1 region with a docking energy of −10.57 kcal/mol. The docked complex was stabilized by three hydrogen bonds between S protein and nelfinavir. E1017, Q954 and Q956 formed hydrogen binding interaction with nelfinavir, whereas I1013 and L1012 formed hydrophobic interactions (Fig. 6B).

**Figure 6.**
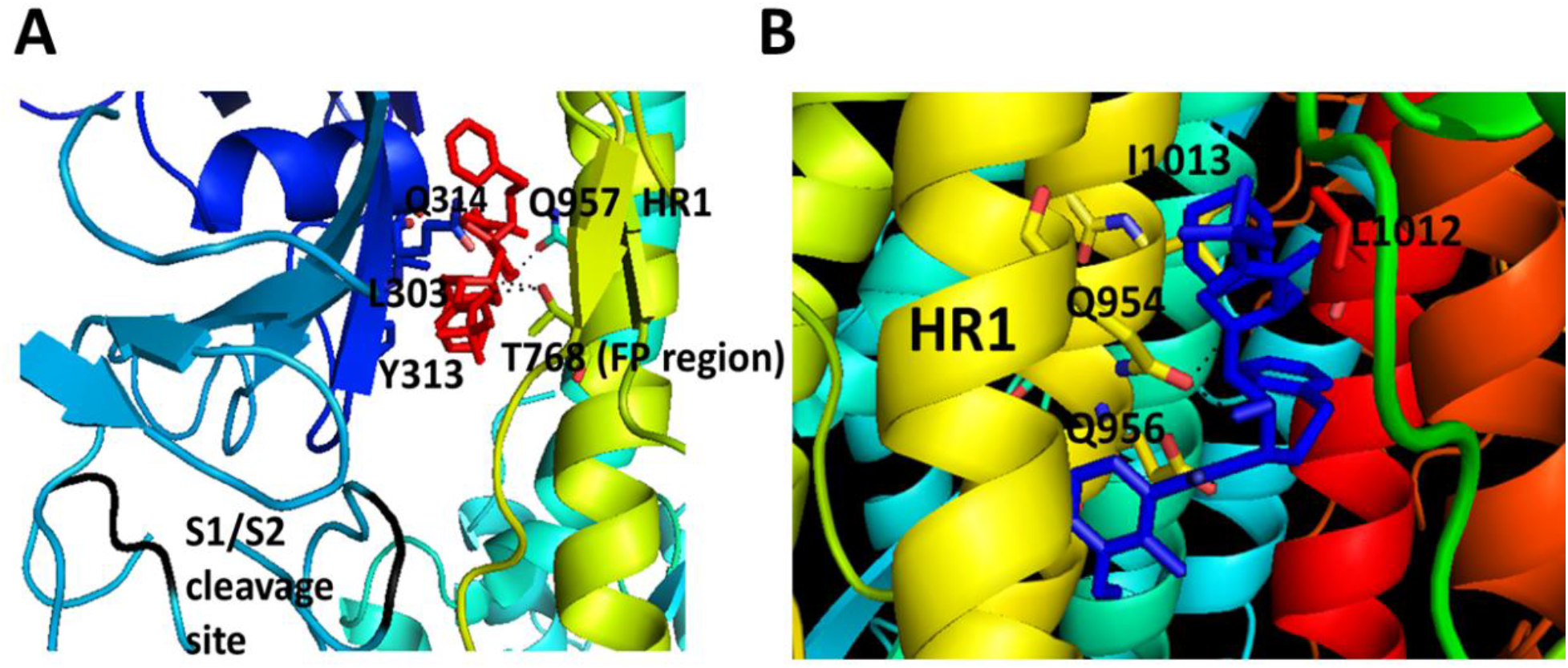
Proposed model of nelfinavir-SARS-2 CoV S protein interaction based on docking. A) Docking near S1/S2 site resulted in docking of nelfinavir binding to fusion peptide (FP) (yellow helical region) and heptad region 1 (HR1) (cyan helix). Nelfinavir is shown in red sticks. Amino acids from the protein that form interaction with nelfinavir are shown as sticks with single letter code for amino acids. B) Docking of nelfinavir on HR1 region. Nelfinavir is shown as blue sticks for the sake of clarity. Amino acids from the protein that form interaction with nelfinavir are shown as blue sticks with single letter code for amino acids.

## Discussion

Virus-induced cell fusion and the formation of multinucleated cells (syncytia) is the hallmark of many different viral infections including retroviruses, herpesviruses, coronaviruses and other viruses. This membrane fusion phenomena are caused by expression of fusogenic glycoproteins on infected cell surfaces. Cell-to-cell fusion mediated by viral glycoproteins is similar to fusion of viral envelopes with cellular membranes that typically occur at the plasma membrane at physiological pH or after endocytosis of virion enveloped particles within endosomes followed by fusion of the viral envelope with endosomal membranes to release the nucleocapsid protein in the cytoplasm (38). Virus-induced cell fusion is an important cytopathic phenomenon because the virus can spread from cell-to-cell avoiding extracellular spaces and exposure to neutralizing antibodies (39). Virus-induced cell fusion can also cause hyper-inflammatory responses producing adverse effects in the infected host. Herein, we show that the SARS CoV-2 Spike (S-n) glycoprotein causes significantly more cell fusion and syncytia formation in comparison to the SARS Spike (S-o) glycoprotein following transient expression in Vero cells. Importantly, we show that nelfinavir mesylate, a currently prescribed anti-HIV protease inhibitor, drastically inhibited both S-n and S-o-mediated cell fusion. These results indicate that it is highly likely that increased SARS CoV-2 virulence over SARS may be attributed to the enhanced fusogenicity exhibited by S-n in comparison to the S-o glycoprotein. Importantly, the fact the nelfinavir significantly inhibited S-n and S-o-mediated cell fusion suggests that it should be tested as an anti-SARS-CoV-2 antiviral, especially at early times after first symptoms are exhibited in infected individuals.

Transient expression of S-n and S-o glycoproteins produced drastic differences in cell fusion, while their overall protein expression was similar, as evidenced by immunohistochemistry signals obtained at 48 hpt. The enhanced fusogenicity of SARS CoV-2 versus SARS was recently noted in infection of Vero cells (40) further validating that our transient transfection results reflect Spike-mediated virus-induced cell fusion differences between SARS and SARS CoV-2. Cell-surface expression of S-n and S-o was comparable suggesting that the observed differences in membrane fusion was due to inherent differences in the structure and function of S-n versus S-o glycoproteins. Interestingly, independent expression of S-n S1 and S2 domains did not cause any cell fusion. It is not clear whether these two domains could be processed and expressed on cell-surfaces, although the S2 domain could be detected via immunohistochemistry (not shown). These results suggest that the entire S glycoprotein needs to be expressed in an uncleaved form that may be proteolytically processed either within endosomes or at cell surfaces by proteases such as TMPRSS2, which is known to be required for Spike activation during virus entry (12).

We utilized the S-n and S-o transient expression system to screen for currently available drugs that may inhibit S-mediated cell fusion and the formation of syncytia. We found that nelfinavir mesylate, a known and currently prescribed anti-HIV drug significantly inhibited S-n and S-o-mediated cell fusion at micromolar concentrations. These results are significant because nelfinavir did not appear to inhibit overall S-n or S-o synthesis and cell-surface expression. Computational modeling revealed that nelfinavir may directly bind to the trimeric form of S-n and S-o near the putative fusogenic domain, and thus, it may directly inhibit S-mediated cell fusion. However, nelfinavir has pleotropic effects on multiple cellular processes including inducing apoptosis an ER stress under certain conditions and has been investigated for anti-cancer purposes (31, 41, 42).

Thus, it is possible that cellular signaling processes are affected that alter the post-translational processing of S-n and S-o, without affecting their cell-surface expression. It is also possible that nelfinavir may inhibit cellular proteases including TMPRSS2 that may be required for S-n and S-o fusion activation. Preliminary experiments indicate that S-n and S-o may be cleaved in Vero cells in the presence of nelfinavir, although it is not currently known whether this cleavage occurs efficiently. In addition, transfected cells expressing S-n or S-o in the presence of nelfinavir did not appear to exhibit morphologies indicating of cellular cytotoxicity, suggesting that nelfinavir is not cytotoxic at the concentrations used in this study. Overall, these experiments suggest that nelfinavir should be used to combat SARS-CoV-2 infections early during the first symptoms exhibited by infected patients to minimize virus spread and provide sufficient time to infected patients to mount a protective immune response.

## References

1. Perlman S. 2020. Another Decade, Another Coronavirus. N Engl J Med 382:760–762.

2. Wu F, Zhao S, Yu B, Chen Y-M, Wang W, Song Z-G, Hu Y, Tao Z-W, Tian J-H, Pei Y-Y, Yuan M-L, Zhang Y-L, Dai F-H, Liu Y, Wang Q-M, Zheng J-J, Xu L, Holmes EC, Zhang Y-Z. 2020. A new coronavirus associated with human respiratory disease in China. Nature 579:265–269.

3. Zhu N, Zhang D, Wang W, Li X, Yang B, Song J, Zhao X, Huang B, Shi W, Lu R, Niu P, Zhan F, Ma X, Wang D, Xu W, Wu G, Gao GF, Tan W. 2020. A Novel Coronavirus from Patients with Pneumonia in China, 2019. New England Journal of Medicine 382:727–733.

4. Demmler GJ, Ligon BL. 2003. Severe acute respiratory syndrome (SARS): a review of the history, epidemiology, prevention, and concerns for the future. Semin Pediatr Infect Dis 14:240–4.

5. Vijayanand P, Wilkins E, Woodhead M. 2004. Severe acute respiratory syndrome (SARS): a review. Clin Med (Lond) 4:152–60.

6. Al Mutair A, Ambani Z. 2020. Narrative review of Middle East respiratory syndrome coronavirus (MERS-CoV) infection: updates and implications for practice. J Int Med Res 48:300060519858030.

7. Badawi A, Ryoo SG. 2016. Prevalence of comorbidities in the Middle East respiratory syndrome coronavirus (MERS-CoV): a systematic review and meta-analysis. Int J Infect Dis 49:129–33.

8. Ramadan N, Shaib H. 2019. Middle East respiratory syndrome coronavirus (MERS-CoV): A review. Germs 9:35–42.

9. Song Z, Xu Y, Bao L, Zhang L, Yu P, Qu Y, Zhu H, Zhao W, Han Y, Qin C. 2019. From SARS to MERS, Thrusting Coronaviruses into the Spotlight. Viruses 11.

10. Du L, Zhao G, Chan CC, Sun S, Chen M, Liu Z, Guo H, He Y, Zhou Y, Zheng BJ, Jiang S. 2009. Recombinant receptor-binding domain of SARS-CoV spike protein expressed in mammalian, insect and E. coli cells elicits potent neutralizing antibody and protective immunity. Virology 393:144–50.

11. Bosch BJ, Martina BE, Van Der Zee R, Lepault J, Haijema BJ, Versluis C, Heck AJ, De Groot R, Osterhaus AD, Rottier PJ. 2004. Severe acute respiratory syndrome coronavirus (SARS-CoV) infection inhibition using spike protein heptad repeat-derived peptides. Proc Natl Acad Sci U S A 101:8455–60.

12. Hoffmann M, Kleine-Weber H, Krüger N, Müller M, Drosten C, Pöhlmann S. 2020. The novel coronavirus 2019 (2019-nCoV) uses the SARS-coronavirus receptor ACE2 and the cellular protease TMPRSS2 for entry into target cells doi:10.1101/2020.01.31.929042. Cold Spring Harbor Laboratory.

13. Matsuyama S, Nagata N, Shirato K, Kawase M, Takeda M, Taguchi F. 2010. Efficient activation of the severe acute respiratory syndrome coronavirus spike protein by the transmembrane protease TMPRSS2. J Virol 84:12658–64.

14. Shulla A, Heald-Sargent T, Subramanya G, Zhao J, Perlman S, Gallagher T. 2011. A transmembrane serine protease is linked to the severe acute respiratory syndrome coronavirus receptor and activates virus entry. J Virol 85:873–82.

15. Bosch BJ, van der Zee R, de Haan CAM, Rottier PJM. 2003. The coronavirus spike protein is a class I virus fusion protein: structural and functional characterization of the fusion core complex. Journal of virology 77:8801–8811.

16. Hofmann H, Geier M, Marzi A, Krumbiegel M, Peipp M, Fey GH, Gramberg T, Pöhlmann S. 2004. Susceptibility to SARS coronavirus S protein-driven infection correlates with expression of angiotensin converting enzyme 2 and infection can be blocked by soluble receptor. Biochemical and biophysical research communications 319:1216–1221.

17. Simmons G, Gosalia DN, Rennekamp AJ, Reeves JD, Diamond SL, Bates P. 2005. Inhibitors of cathepsin L prevent severe acute respiratory syndrome coronavirus entry. Proceedings of the National Academy of Sciences of the United States of America 102:11876–11881.

18. Tripet B, Howard MW, Jobling M, Holmes RK, Holmes KV, Hodges RS. 2004. Structural characterization of the SARS-coronavirus spike S fusion protein core. J Biol Chem 279:20836–49.

19. Wong SK, Li W, Moore MJ, Choe H, Farzan M. 2003. A 193-Amino Acid Fragment of the SARS Coronavirus S Protein Efficiently Binds Angiotensin-converting Enzyme 2. Journal of Biological Chemistry 279:3197–3201.

20. Yang Z-Y, Huang Y, Ganesh L, Leung K, Kong W-P, Schwartz O, Subbarao K, Nabel GJ. 2004. pH-dependent entry of severe acute respiratory syndrome coronavirus is mediated by the spike glycoprotein and enhanced by dendritic cell transfer through DC-SIGN. Journal of virology 78:5642–5650.

21. Dimitrov DS. 2003. The Secret Life of ACE2 as a Receptor for the SARS Virus. Cell 115:652–653.

22. Xia S, Zhu Y, Liu M, Lan Q, Xu W, Wu Y, Ying T, Liu S, Shi Z, Jiang S, Lu L. 2020. Fusion mechanism of 2019-nCoV and fusion inhibitors targeting HR1 domain in spike protein. Cellular & Molecular Immunology doi:10.1038/s41423-020-0374-2.

23. Wrapp D, Wang N, Corbett KS, Goldsmith JA, Hsieh C-L, Abiona O, Graham BS, McLellan JS. 2020. Cryo-EM structure of the 2019-nCoV spike in the prefusion conformation. Science 367:1260–1263.

24. Pai VB, Nahata MC. 1999. Nelfinavir Mesylate: A Protease Inhibitor. Annals of Pharmacotherapy 33:325–339.

25. Tebas P, Powderly WG. 2000. Nelfinavir mesylate. Expert Opinion on Pharmacotherapy 1:1429–1440.

26. Gupta AK, Li B, Cerniglia GJ, Ahmed MS, Hahn SM, Maity A. 2007. The HIV protease inhibitor nelfinavir downregulates Akt phosphorylation by inhibiting proteasomal activity and inducing the unfolded protein response. Neoplasia 9:271–8.

27. Pajonk F, Himmelsbach J, Riess K, Sommer A, McBride WH. 2002. The human immunodeficiency virus (HIV)-1 protease inhibitor saquinavir inhibits proteasome function and causes apoptosis and radiosensitization in non-HIV-associated human cancer cells. Cancer Res 62:5230–5.

28. Yang Y, Ikezoe T, Takeuchi T, Adachi Y, Ohtsuki Y, Takeuchi S, Koeffler HP, Taguchi H. 2005. HIV-1 protease inhibitor induces growth arrest and apoptosis of human prostate cancer LNCaP cells in vitro and in vivo in conjunction with blockade of androgen receptor STAT3 and AKT signaling. Cancer Science 96:425–433.

29. Bruning A, Burger P, Vogel M, Rahmeh M, Gingelmaiers A, Friese K, Lenhard M, Burges A. 2009. Nelfinavir induces the unfolded protein response in ovarian cancer cells, resulting in ER vacuolization, cell cycle retardation and apoptosis. Cancer Biol Ther 8:226–32.

30. Guan M, Su L, Yuan YC, Li H, Chow WA. 2015. Nelfinavir and nelfinavir analogs block site-2 protease cleavage to inhibit castration-resistant prostate cancer. Sci Rep 5:9698.

31. Pyrko P, Kardosh A, Wang W, Xiong W, Schonthal AH, Chen TC. 2007. HIV-1 protease inhibitors nelfinavir and atazanavir induce malignant glioma death by triggering endoplasmic reticulum stress. Cancer Res 67:10920–8.

32. Yamamoto N, Yang R, Yoshinaka Y, Amari S, Nakano T, Cinatl J, Rabenau H, Doerr HW, Hunsmann G, Otaka A, Tamamura H, Fujii N, Yamamoto N. 2004. HIV protease inhibitor nelfinavir inhibits replication of SARS-associated coronavirus. Biochem Biophys Res Commun 318:719–25.

33. Petit CM, Chouljenko VN, Iyer A, Colgrove R, Farzan M, Knipe DM, Kousoulas KG. 2007. Palmitoylation of the cysteine-rich endodomain of the SARS-coronavirus spike glycoprotein is important for spike-mediated cell fusion. Virology 360:264–74.

34. Petit CM, Melancon JM, Chouljenko VN, Colgrove R, Farzan M, Knipe DM, Kousoulas KG. 2005. Genetic analysis of the SARS-coronavirus spike glycoprotein functional domains involved in cell-surface expression and cell-to-cell fusion. Virology 341:215–30.

35. Morris GM, Huey R, Lindstrom W, Sanner MF, Belew RK, Goodsell DS, Olson AJ. 2009. AutoDock4 and AutoDockTools4: Automated docking with selective receptor flexibility. J Comput Chem 30:2785–91.

36. Kozisek M, Bray J, Rezacova P, Saskova K, Brynda J, Pokorna J, Mammano F, Rulisek L, Konvalinka J. 2007. Molecular analysis of the HIV-1 resistance development: enzymatic activities, crystal structures, and thermodynamics of nelfinavir-resistant HIV protease mutants. J Mol Biol 374:1005–16.

37. Wrapp D, Wang N, Corbett KS, Goldsmith JA, Hsieh CL, Abiona O, Graham BS, McLellan JS. 2020. Cryo-EM structure of the 2019-nCoV spike in the prefusion conformation. Science 367:1260–1263.

38. Podbilewicz B. 2014. Virus and cell fusion mechanisms. Annu Rev Cell Dev Biol 30:111–39.

39. Mothes W, Sherer NM, Jin J, Zhong P. 2010. Virus cell-to-cell transmission. J Virol 84:8360–8.

40. Xia S, Liu M, Wang C, Xu W, Lan Q, Feng S, Qi F, Bao L, Du L, Liu S, Qin C, Sun F, Shi Z, Zhu Y, Jiang S, Lu L. 2020. Inhibition of SARS-CoV-2 (previously 2019-nCoV) infection by a highly potent pan-coronavirus fusion inhibitor targeting its spike protein that harbors a high capacity to mediate membrane fusion. Cell Res 30:343–355.

41. Gills JJ, LoPiccolo J, Dennis PA. 2008. Nelfinavir, a new anti-cancer drug with pleiotropic effects and many paths to autophagy. Autophagy 4:107–109.

42. Gills JJ, LoPiccolo J, Tsurutani J, Shoemaker RH, Best CJM, Abu-Asab MS, Borojerdi J, Warfel NA, Gardner ER, Danish M, Hollander MC, Kawabata S, Tsokos M, Figg WD, Steeg PS, Dennis PA. 2007. Nelfinavir, A Lead HIV Protease Inhibitor, Is a BroadSpectrum, Anticancer Agent that Induces Endoplasmic Reticulum Stress, Autophagy, and Apoptosis In vitro and In vivo. Clinical Cancer Research 13:5183–5194.

