## Supplemental Figure 1 for "The anti-HIV Drug Nelfinavir Mesylate (Viracept) is a Potent Inhibitor of Cell Fusion Caused by the SARS-CoV-2 Spike (S) Glycoprotein Warranting further Evaluation as an Antiviral against COVID-19 infections"

**Supplementary Figure 1s**

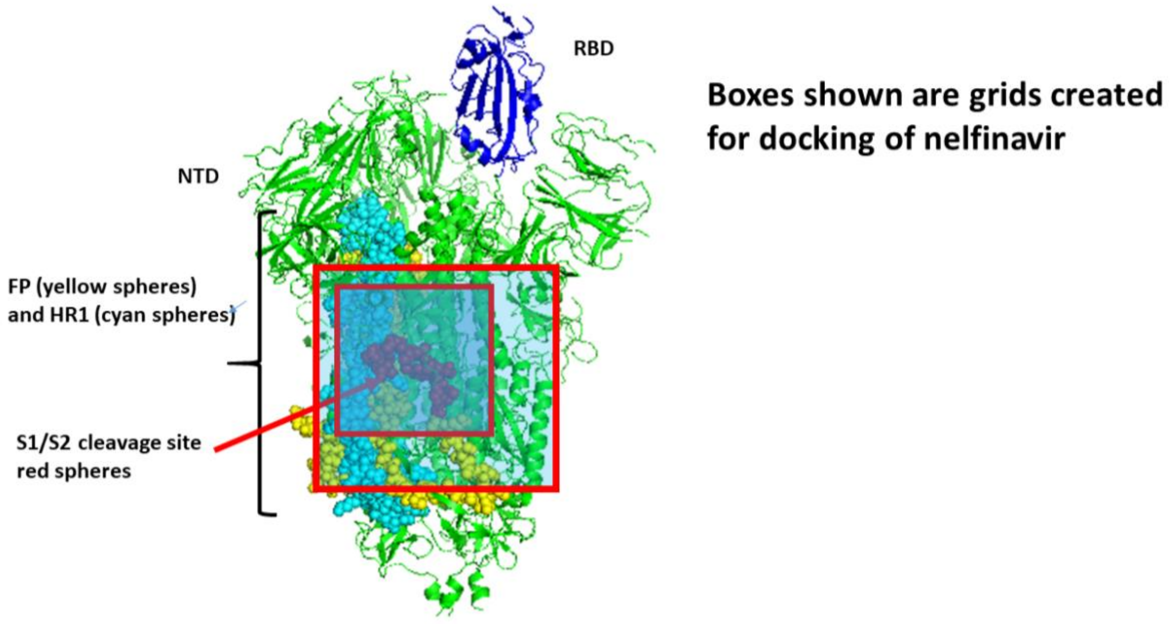


**Figure. S1**. Structure of S protein of SARS CoV-2 with different regions labeled. Grids used for docking calculations are shown as boxes. NTD-N-terminal domain, RBD-receptor binding domain, FP-fusion peptide, HR1-heptad region-1.
